# Biodiversity effects on seedling biomass growth are modulated by light environment across functional groups

**DOI:** 10.1101/2022.03.08.483461

**Authors:** Krishna Anujan, Alisha Shabnam, Irfan Ali, G Ashok Kumar, Mahesh Sankaran, Meghna Krishnadas, Shahid Naeem

## Abstract

1. Tree biodiversity has the potential to ensure consistency in the functioning of forest ecosystems, not just over space, but over long-timescales by maintaining composition through recruitment. However, for continued buffering in the face of global environmental change, the sensitivity of biodiversity-ecosystem functioning relationships to heterogeneous environments needs to be understood.
2. Seedling recruitment in carbon-rich tropical forests is a result of biotic and abiotic drivers but their combined outcomes at the community-level remain poorly understood. Although biodiversity in seedling communities can potentially increase their growth and biomass accumulation, abiotic drivers like light can alter this effect through divergent effects on constituent species and functional groups. In forests with high baseline heterogeneity in microclimates, these processes can enhance or constrain regeneration.
3. We tested the effects and interactions between species richness and canopy cover on the growth of seedling communities consisting of tropical broad-leaved evergreen and deciduous forest species using a fully crossed manipulated experiment in the Andaman Islands, India and compared these with field observations from a long-term forest plot in the same landscape.
4. We show that in the critical seedling establishment phase, species richness and light increase community biomass independently. Accounting for variation across species, individual species on average accumulated more biomass in communities with both higher light and higher diversity.
5. We also show that overyielding in species rich communities fits expectations from a model of complementarity with non-random overyielding than selection or spatial insurance effects.
6. *Synthesis* Taken together, our results show that the potential for biodiversity to increase ecosystem functioning in seedling communities is modulated by light. Further understanding on the interaction of biodiversity with multiple abiotic drivers and their effect on regeneration dynamics is crucial for predicting future ecosystem functioning.

## Introduction

Diverse forests consistently store more carbon than low diversity stands, but these ecosystem functions can be destabilised by changing environments is less understood (Ammer, 2019; Hutchison et al., 2018). At global and regional scales, higher tree diversity in forests is associated with greater productivity and higher contribution towards reducing atmospheric carbon (Liang et al., 2016; Osuri et al., 2020). However, with global environmental change increasing spatial and temporal environmental heterogeneity and driving compositional shifts, the future of forest-associated carbon storage remains uncertain (Feeley et al., 2011). In forested ecosystems, where large, long-lived trees have disproportionate contributions to biomass, productivity and associated ecosystem functions, future functioning depends on mortality and recruitment processes that shape community composition (Bunker et al., 2005). Although the responses of tree mortality to global environmental change are actively studied, recruitment processes have received relatively less attention in this context (but see Zhao et al., 2018). Understanding the importance of diversity to the community function of recruited tree communities under heterogenous environments is crucial for understanding and predicting the future of ecosystem functioning under scenarios of co-occurring environmental change and biodiversity loss.

The contribution of species diversity to ecosystem functioning under varied environmental conditions depends on how these conditions affect species performance. Diverse communities could have higher yield compared to monocultures either due to increased probability of selecting species that are high yielding across all conditions (selection effect), higher yield across all species in mixtures across all conditions (complementarity), each species performing better in a suitable condition (spatial insurance), or differences in species yield but species overyielding in mixtures (complementarity with non-random overyielding) (Isbell et al., 2018). Trait differences between species and their performance under different conditions can therefore contribute significantly to the biodiversity effect, with larger spatial heterogeneity requiring higher species richness to maintain biodiversity (Thompson et al., 2018, 2021). Tests of these basic theoretical frameworks have mostly involved grassland communities with annuals or short-lived perennials. On the other hand, tropical forests store roughly 55% of terrestrial carbon with 56% of this stored in live biomass (Badgley et al., 2019; Pan et al., 2011). In these forests with long-lived species, the relative importance of diversity mechanisms, fine-scale environmental heterogeneity and high species richness remains unknown.

Intraspecific competition plays an integral role in structuring seedling communities and neighbourhood biodiversity can therefore increase growth. Studies with tropical tree species show that species richness and functional diversity in communities has the potential to increase the relative growth rate of individual seedlings and cause overyielding in communities (Kuptz et al., 2010; Sapijanskas et al., 2013; Shen et al., 2021; Van de Peer et al., 2018). Seedling diversity in the neighbourhood can alter seedling root traits directly, altering resource niches and acquisition patterns, and increasing growth at the community level (Madsen et al., 2020). However, seedling competition occurs in an environmental context for limiting resources, and the outcomes of interspecific interactions can change as the environment varies (Butterfield & Callaway, 2013). Biodiversity can therefore increase spatial and temporal insurance, leading to more consistent growth (lower variation over time), and higher biomass accumulation compared to single-species stands (Hutchison et al., 2018; Isbell et al., 2018; Tuck et al., 2016).

In diverse forests, light differentially affects growth across functional groups of tree seedlings, potentially structuring local communities and affecting standing biomass along natural gradients. As a result of pervasive light competition in forest understories, seedlings respond to increased light intensity with increased growth (Kuptz et al., 2010; Lu et al., 2021; Sangsupan et al., 2021; Sovu et al., 2010). Fundamentally, growth plasticity and the ability to utilize increased light is mediated by species identity or functional identity (Kuptz et al., 2010; Tomlinson et al., 2014; Tripathi et al., 2020). Deciduous species grow faster under increased light conditions while evergreen species have slower growth and are sensitive to drought in high light conditions (Tripathi et al., 2020). Mixed deciduous forests, a type of seasonally dry tropical forests, have both broad-leaved evergreen and deciduous species in the same canopy, leading to seasonal heterogeneity of microclimates in the forest floor. These differences can determine the distribution of seedlings in the understory; deciduous species in the canopy increase the abundance and diversity of light-demanding species in the understory by increasing light availability (Souza et al., 2014). Since evergreen and deciduous species are adapted to low and high light respectively, potential diversity effects under different light conditions are likely driven by mechanisms of spatial insurance. However, with increased tree mortality in tropical forests due to drought and extreme climatic events (Aleixo et al., 2019; Uriarte et al., 2019), light environments in the understory are potentially undergoing large-scale shifts, affecting both biodiversity and its effects on ecosystem functioning.

We experimentally assessed the combined influence of light and species diversity on the growth of tropical forest seedling communities and compared dynamics with field communities under natural regeneration in a high diversity tropical forest. We expected that (i) light would increase overall seedling growth and lead to overyielding in high light treatments compared to shade treatments (ii) decreased intraspecific competition in mixed cultures would lead to higher biomass with increasing species richness and combined with expectation (i), lead to higher overyielding in high diversity, high light treatments (iii) light responses of evergreen and deciduous species would be different, with evergreen species performing better under low light and deciduous species performing better under high light environments, i.e., overyielding through spatial insurance.

## Methods

We measured the combined influences of two key factors structuring forests – light and species diversity – through experimental methods and long-term data from a tropical forest landscape. The study was conducted in the Andaman Islands, India, an archipelago in the Bay of Bengal. It is part of the Indo-Burma biodiversity hotspot with high species diversity, high endemism and >80% forest cover. Many forests in the archipelago are mixed deciduous forests, where tropical evergreen and tropical deciduous tree species co-exist in the canopy. This has the potential to create heterogeneity in microclimate in the understory. A shadehouse experiment was conducted at the Silviculture Research nursery, Nayashahar, maintained by the Department of Environment and Forest, Andaman and Nicobar Islands. We matched the design of the experimental plots (plot size, range of species richness, abundance) and census methods closely with seedling plots at a permanent forest dynamics plot at Alexandra Island, about 10 km from the experiment site and with mixed deciduous forest type. This plot is part of the Long-term Ecosystem Monitoring Network (LEMoN) in India established to set baselines for forest dynamics and carbon storage across various biomes in India.

### Experimental Methods

In a fully crossed experiment, we planted similarly sized evergreen and deciduous seedlings under 3 shade conditions (simulating an evergreen canopy that is closed throughout the year – “closed”, a deciduous canopy that is closed in the wet season and open in the dry season - “deciduous” and a consistently open condition – “open”) and in 3 levels of diversity (1 species, 3 species and 6 species from a pool of 10 species) and measured growth for 1 year (Fig 1). Each treatment had 6 replicates each, totalling 54 plots. Each plot of 1 m x 1 m plot was planted with 18 individually tagged seedlings belonging to 1, 3 or 6 species from a pool of 10 species in March 2019. Seedling plots of 1m^2^ in the landscape contained an average of 16 seedlings and had a maximum of 7 species and the planting numbers were matched to these naturally occurring values. Experimental plots were set up using natural, well-mixed soil from an abandoned plot nearby and demarcated as regular rows using bamboo and other locally available building material. Over each plot, standard shade nets of 50% shade were used to cover the canopy at 6ft height to mimic either a closed canopy (closed throughout the year) or a deciduous canopy (closed in the wet season and open in the dry season) or they were left fully open throughout the year. The plots were arranged in regular rows with a spacing of 30 cm between adjacent plots and the 9 treatments were randomly distributed spatially among the 54 plots in the experiment.

**Figure 1:**
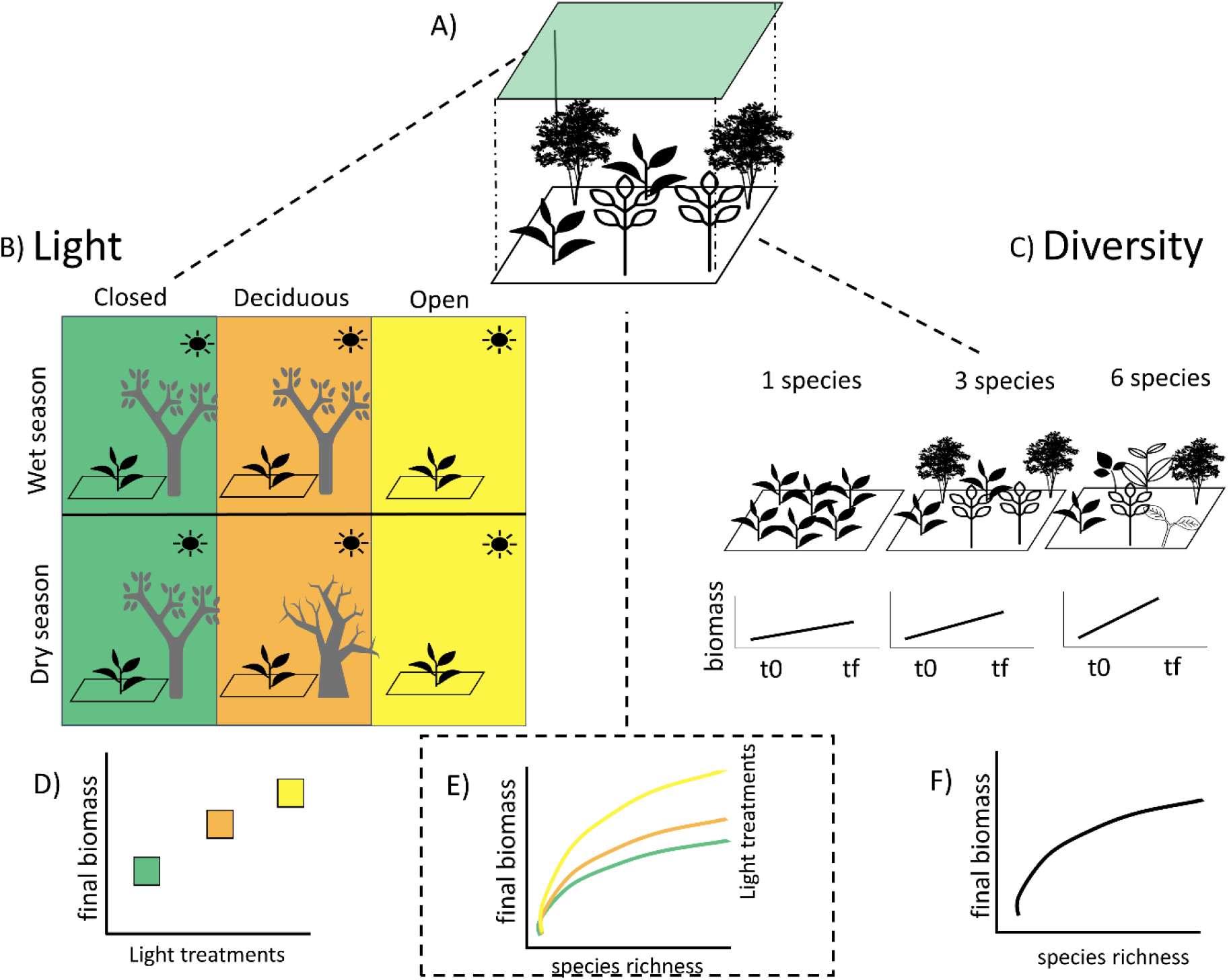
Schematic of experimental and expectations A) Experimental setup with fully crossed light and diversity treatments B) Light treatments in detail – closed, deciduous and open canopy treatments and with colour codes – in the wet and dry season and D) final biomass expectations for each of these based on increased growth in the presence of light. C) Three levels of diversity treatments – 1, 3 and 6 species and biomass expectations between initial and final communities and F) expectations for final biomass between the three treatments – an increasing, saturating curve. And finally, E) combined expectations for community biomass under the combined influence of light and diversity treatments; the influence of species diversity on growth is expected to be modulated by light conditions. Icons from The Noun Project.

We used seedlings of ten native forest tree species germinated by ANI forest department nurseries from seeds collected from nearby forests and for the purpose of replanting into selectively logged forests in the area. Six common species – 3 evergreen (*Dipterocarpus griffithii*, *Myristica andamanica* and *Mangifera andamanica*) and 3 deciduous (*Terminalia procera*, *Pterocarpus dalbergioides* and *Planchonia valida*) were chosen as the primary experimental species to make up the monocultures, while the other 4 species (*Artocarpus chaplasha*, *Diospyros sp.*, *Planchonella longipetiolata*, *Walsura* sp.) were planted in a few of the 3 and 6 species plots, to increase variation in species combinations in polycultures. For the 6 primary species, monocultures were planted under each of the 3 canopy treatments. Seedlings were planted into the experiment when approximately 30 cm tall and within the height range in which these are usually replanted into forests. During transplanting, we removed the plants from the potting soil and measured root length, shoot length, basal area and total biomass, allowing us to create allometric equations of this life history stage. Eighteen individually tagged seedlings were planted into each plot in a random order but with equal spacing, 15 cm apart. After transplanting, we watered the plots for two weeks to reduce mortality specifically from transplanting. Other than weeding every month to maintain planting conditions, the plots were left unmanaged until the end of the experiment.

Two months after replanting, we initiated monthly censuses of the plots for 11 iterations. During each census, the height and basal area of every tagged individual seedling was measured. Height of the seedlings were measured using a standard measuring tape (carpentry tape) with an accuracy of 1 mm. Basal area of seedlings was measured using an Insize digital calliper with an accuracy of 0.01 mm. We report values from the final census, conducted in June 2020 throughout this study.

### Observational data

Seedling data from long-term plots with comparable species composition and microclimate heterogeneity in the landscape were analysed to validate experimental results. Under the LEMoN network, a 1-hectare long-term monitoring plot was set up at Alexandra Island in the Andaman archipelago. This site is in a mixed deciduous forest with no recent logging history and protected as part of the Mahatma Gandhi Marine National Park. Within the 1 ha plot, 25 seedling plots of 1 m^2^ have been set up on a uniform grid covering the extent of the plot. Within these subplots, naturally regenerating seedlings are tagged, identified to the best possible degree and measured at monthly intervals. For comparing the light environments of the 25 different seedling plots, the canopy cover at each location was measured in December 2020, using a Forestry Suppliers Spherical Crown Densiometer, Convex Model A. At each seedling plot, the percent of canopy cover was calculated as the number of grid cells in the densiometer with canopy cover as an average over four replicate measures. Since much of the canopy cover in these plots is determined by the upper canopy where dynamics are slow, these light environments were assumed to be stable through the years of measurements.

Data on basal diameter and height of individual seedlings in these plots measured using similar equipment and methods as the experiment from three censuses done in June 2016, 2017 and 2018 were used for comparison. Height was measured for all individuals in the seedling plot, while basal diameter was measured only for seedlings with height > 1m. For those instances where basal area was not recorded as part of monitoring protocol, basal area was calculated as a function of height using a fit for basal area to height from experimental communities. The allometric equation was *basal diameter* = 2.34 + 0.08 ∗ *height*. Being under natural regeneration, the abundance, species richness and species composition varied across these plots. Over the three censuses, 585 seedlings were recorded, out of which 310 were unique individuals, while the rest were repeated encounters. Gaps in species IDs within the data were filled to the best possible extent using other information in the data sheet – like local names and remarks, but unidentified individuals with no further information were considered as a single species to avoid overcounting of richness. 254 of these individuals remained unassigned to species. Identified species belonged to genera that were used in the manipulated experiment, including *Myristica spp.*, *Terminalia bialata*, *Diospyros spp.*, *Dipterocarpus spp.*, and *Planchonia andamanica*.

### Statistical methods

All statistical analyses were performed in R with R Studio (R Core Team 2020). For both experimental communities and field communities, we analysed the effects of experimental treatments of light and species richness on basal area and height using generalized linear models using the *lme4* package. To avoid the confounding effects of differential survival on total final biomass, we modelled treatment effects on the mean values of surviving individuals in the experiment (n=651) and on observed individuals in the field. Mean basal area and height being continuous, non-zero variables were modelled with gamma errors for continuous, non-zero variables, and because it best fit the distribution.

For the final communities, we assessed the degree of overyielding with species richness and the relative contribution of complementarity and selection effects using methodology from Loreau M. & Hector. A., 2001. We used data from the tenth census of the experiment as one of the monocultures (of *D. griffithii*) had all died by the last census. We used 49 of 54 replicates for this analysis, excluding mixed communities planted with species for which monocultures were not planted in the experiment (see above section). For each species, we calculated the average basal area in a monoculture across all three light treatments (*M*). We calculated relative yield (*ΔY*) in mixed communities as the difference of the observed yield (*Y*) from the expected simple sum. Further, for each plot, we partitioned this difference into selection effects (*cov*(*ΔRY*, *M*), where *ΔRY* = *Y*/_*M*_ − 1/_*N*_) and complementarity effects 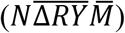, where *N* is the planted species diversity.

For observed communities on field, we modelled total basal area and height of individuals as a generalized linear mixed effects model with a gamma family of errors. We modelled each of these with separate quadratic models of species richness and canopy openness, with intercepts allowed to vary with the year of observation. We compared models with the linear only model (without the quadratic term) using AIC values.

All figures were created using *ggplot2* and *sjPlot*.

## Results

### Growth at the community level

Seedlings in experimental communities with greater species richness and greater light levels accumulated more biomass on average. Individual wet weight at the initial time point was more strongly associated with basal diameter than height (linear model for weight adjusted R^2^ with height=0.26, basal diameter=0.40). Mean basal area in the plant community, indicative of biomass, significantly increased with increasing levels of light and species richness; the 6 species treatment and the deciduous canopy treatment had significantly higher mean basal area than the closed, monocultures (Fig 2). Mean height of the seedlings, however, was only influenced by light; the deciduous treatment significantly increased mean height, while other treatments were not significantly different from the closed canopy monocultures (Fig 2). Moreover, there were no significant interaction effects of light and species richness treatments on either mean basal area or mean height of the seedlings (Fig 2). The differences between communities also persisted through time when analysed at the monthly timescale (repeated measures ANOVA, Fig S2). More trivially, regardless of treatment, higher values of final abundance were strongly associated with higher mean basal area and mean height in these communities (Fig 2). Summed across all individuals in a 1m^2^ plot, total height and basal area in final communities were highly correlated (Pearson’s correlation coefficient r=0.96). The survival of seedlings in plots was not significantly predicted by either planted species richness or by canopy treatment, but communities with higher proportion of evergreen species had significantly lower survival at the end of the experiment (Fig S1).

**Figure 2:**
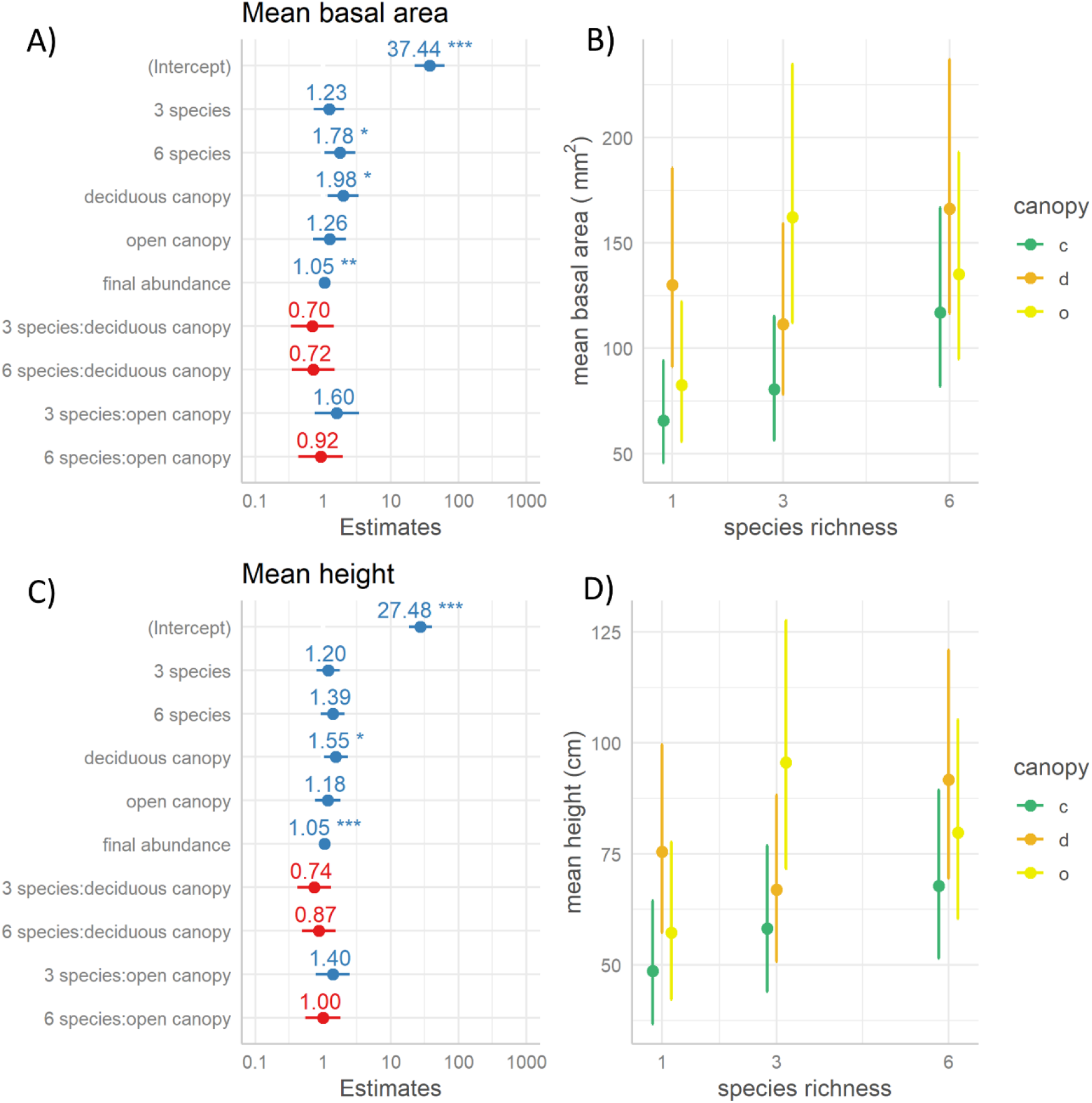
Generalized linear model of mean basal area and mean height of a seedling in a 1m x 1m plot in the experiment. For mean basal area, deviance is 12.96, df=47, p=0.999, R2=0.41. For mean height, deviance =8.24, df=47, p=1, R2=0.48. For mean basal and mean height in a community A) and C) show model estimates of the parameters, B) and D) show model predictions for species richness and canopy treatments. Colour codes for canopy treatment, c=closed, d=deciduous and o=open treatments as specified in Fig1.

Species richness in communities increased overyielding and complementarity effect and decreased the magnitude of selection effect across light environments (Fig 3). At the end of 10 months of the experiment, observed total basal area in many plots was higher than from the expectation from monoculture averages; relative yield was largely positive. The relative yield increased with both light and diversity treatments; two-way ANOVA of observed – expected total basal area was significant for both species richness treatments (F=13.6, df=1, p<0.001***) and canopy treatments (F=3.4, df=2, p=0.04*). Further, the effect of complementarity increased and selection effect decreased with increasing species richness (Fig 3). At the highest level of species diversity, the six species treatment, the magnitude of complementarity effect across all light treatments in all communities were positive while selection effect was negative for all communities. The magnitude of the complementarity effect was significantly different across light and diversity treatments (two-way ANOVA: species richness F=24.23, df=1, p<0.001***, canopy treatment F=3.74, df=2, p=0.03*), while selection effect was significantly different only with species richness (F=23.3, df=1, p<0.001***, canopy treatment F=1.121, df=2, p=0.33).

**Figure 3:**
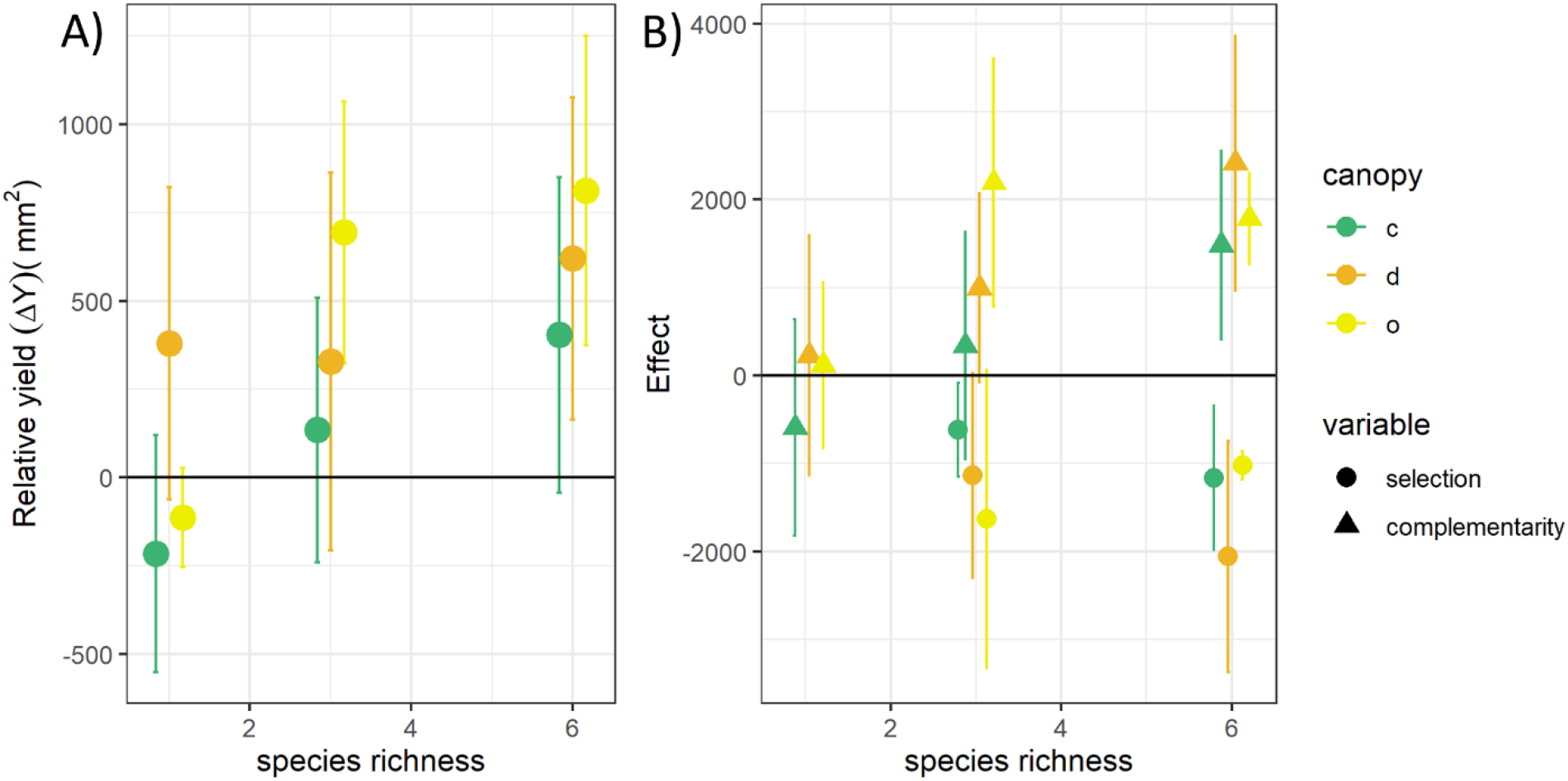
Relative contributions of complementarity and selection effect to relative yield (*ΔY*) of multispecies communities. A) Plot of the relative yield (*ΔY*) for different treatments, based on basal area measurements. B) The magnitude of complementarity and selection effects, as calculated by the Loreau-Hector method (details in methods). Points represent means and errors bars represent 95% confidence intervals.

### Predictors of growth at the individual level

Accounting for differences in species identity, individual seedlings that survived until the end of the experiment showed higher gain in basal diameter and height in mixed communities and in the deciduous treatment as compared to counterparts in closed monocultures (n=651, RMSE=3.21, marginal R2=0.11; Fig 4). However, the open canopy treatment was not significantly different from the closed treatment. Models also revealed significant negative interactions between the species richness and light treatments for individual seedling growth (Fig 4). Although species richness increased basal diameter and height across light treatments, this gain was lower in the deciduous treatment than in the closed treatment. Basal area and height at planting significantly increased corresponding gains in these response variables within a year of the experiment (Fig 4).

**Figure 4:**
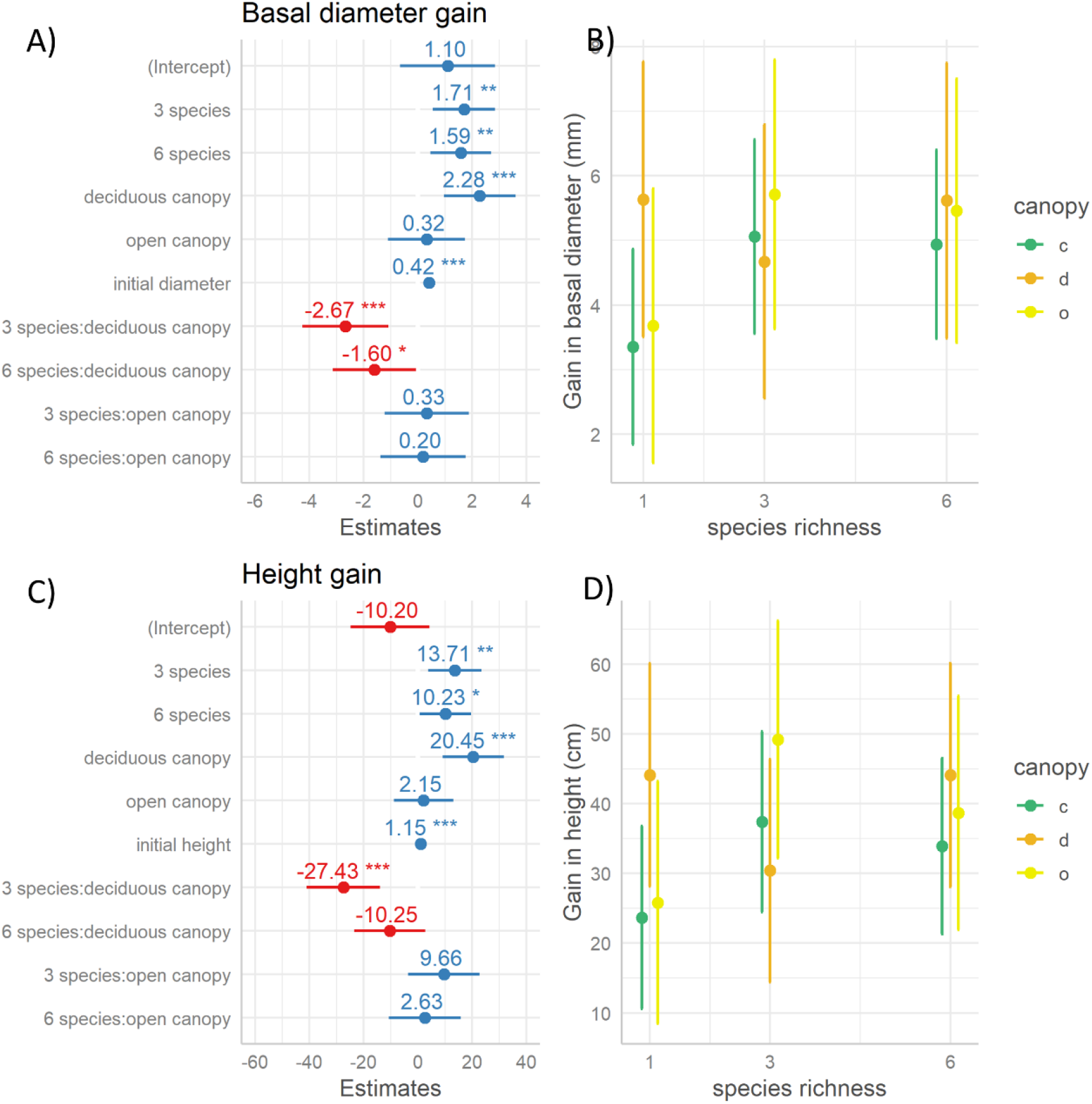
Linear mixed effects models for gain in basal diameter and height between the first and the last census for individual seedlings; random slope model with variable slopes for each species with canopy treatment. For basal diameter, deviance=3414, total R^2^=0.44, fixed effect R^2^=0.16; for height, deviance=6200, total R^2^=0.50, fixed effect R^2^=0.29. For basal diameter and height respectively, A) and C) represent estimated effects of different predictors, B) and D) show predicted responses from the model for species richness and canopy conditions

Individual growth varied with species in the experiment and under different canopy conditions, but the differences remained consistent across light treatments. The model with species specific slopes, varying with canopy, had the lowest AIC, lower than the random intercept model (ΔAIC (for df=17 and df=12) =2.05). All other models - with no random effects and with only random effects – performed much worse. The response of biomass to species diversity and light dependent on species identity; in the full model with species richness, canopy treatment and slopes varying with species, the random effect of species on slopes was significant (SD of random effect species identity at closed canopy = 1.89; deciduous canopy = 1.09, open canopy = 1.20, residual =3.26). Deciduous species – *T. procera*, *P. valida*, *P. dalbergioides* and the evergreen *D. griffithii* had higher than average growth rates, demonstrated by positive values of the random effect, while *Myristica andamanica* and *Mangifera andamanica* were slower growing species (Fig 5). These patterns were consistent between basal diameter and height. Moreover, these differences were consistent across light treatments, although starker under lower light.

**Figure 5:**
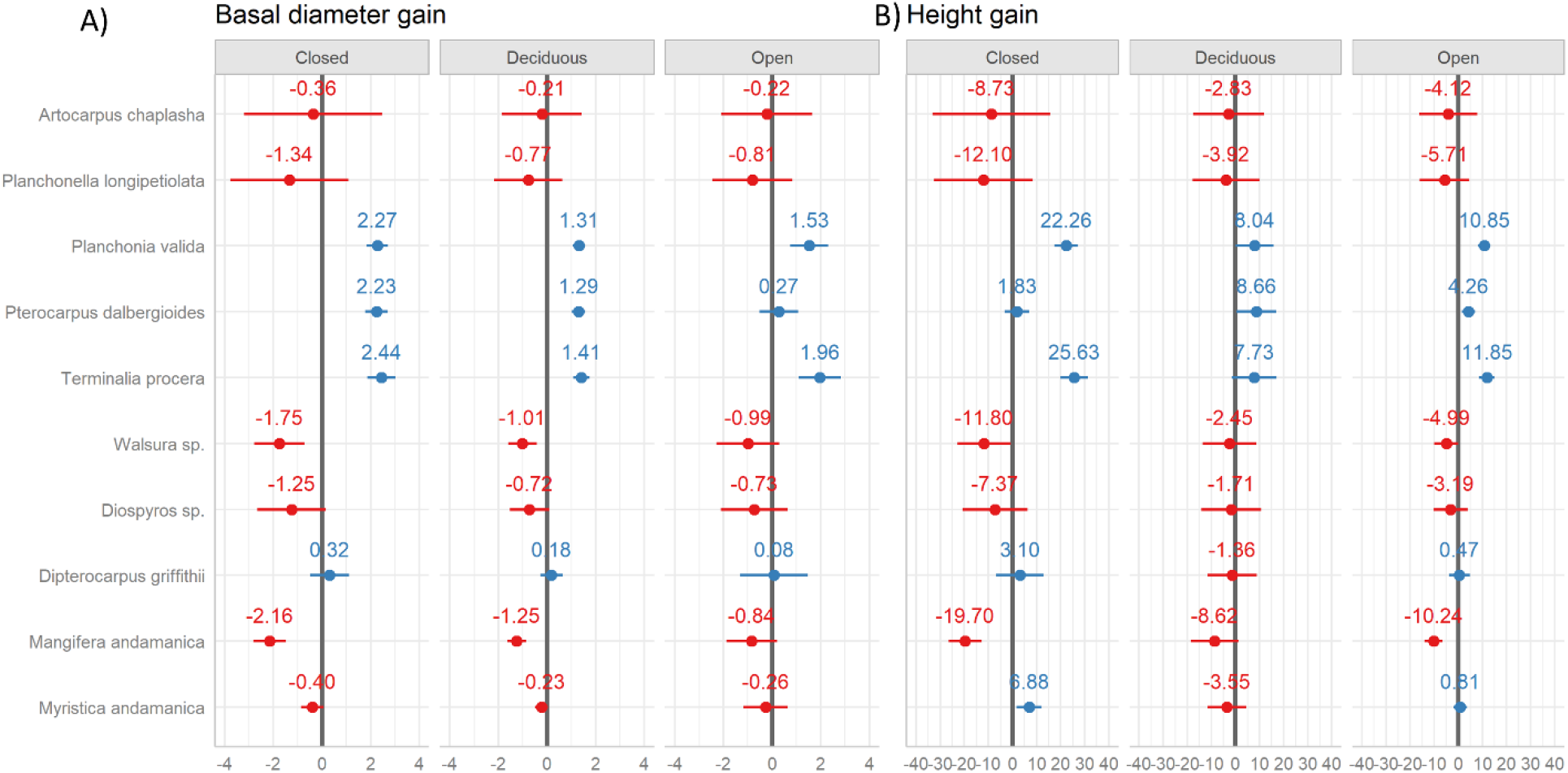
Random slopes of species under different canopy conditions; A) shows random effects for each species for gain in basal diameter and B) for height gain. The top six species represent fast-growing, deciduous species, followed by slower growing, evergreen species towards the bottom.

Species functional identities affect total basal area and total height of the community. Although functional diversity levels were not fully replicated in the experimental design, the proportion of evergreen species in the community affected the final biomass across all experimental treatments. Overall, increased proportion of evergreen species was associated with lower total basal area and total height across different light treatments (Fig S3). In the open canopy treatment, however, a 0.3 proportional abundance of evergreen species had the highest total basal area and total height. Due to a lack of replication of this combination in the closed and deciduous treatments, further tests of this pattern are not possible.

### Convergence with observational data

Higher values of biomass accumulation in seedling communities in observational plots was associated with higher species richness and canopy openness. For 25 plots across a range of light conditions and species richness, separate models of total basal area and height in seedling communities showed significant non-linear effects of canopy openness and species richness on these variables. AIC values of all quadratic models were lower than for models with linear terms alone; except for height and species richness, where there was no clear best-fit model (Table S1).

Total basal area in communities increased with species richness, but peaked at intermediate levels; for a model with observation year as a random effect, the positive linear and negative quadratic term were significant (RMSE=0.5, marginal R^2^=0.99, Fig 6A). Total basal area had a quadratic relationship with canopy openness (RMSE=0.50, marginal R^2^=0.89, Fig 6B). Similarly, total height of seedling communities was highest at intermediate openness; a quadratic model of canopy openness was significant (RMSE=0.49, marginal R^2^=0.94, conditional R^2^=0.99, Fig 6D). However, a quadratic model of total height with species richness only showed marginal effect of the linear term of species richness (RMSE=0.51, marginal R^2^=0.99. Fig 6C). Across all treatments, the values of total basal area and total height in the field observations were lower than the experimental communities, but total height was closer in magnitude (Fig 6).

**Figure 6:**
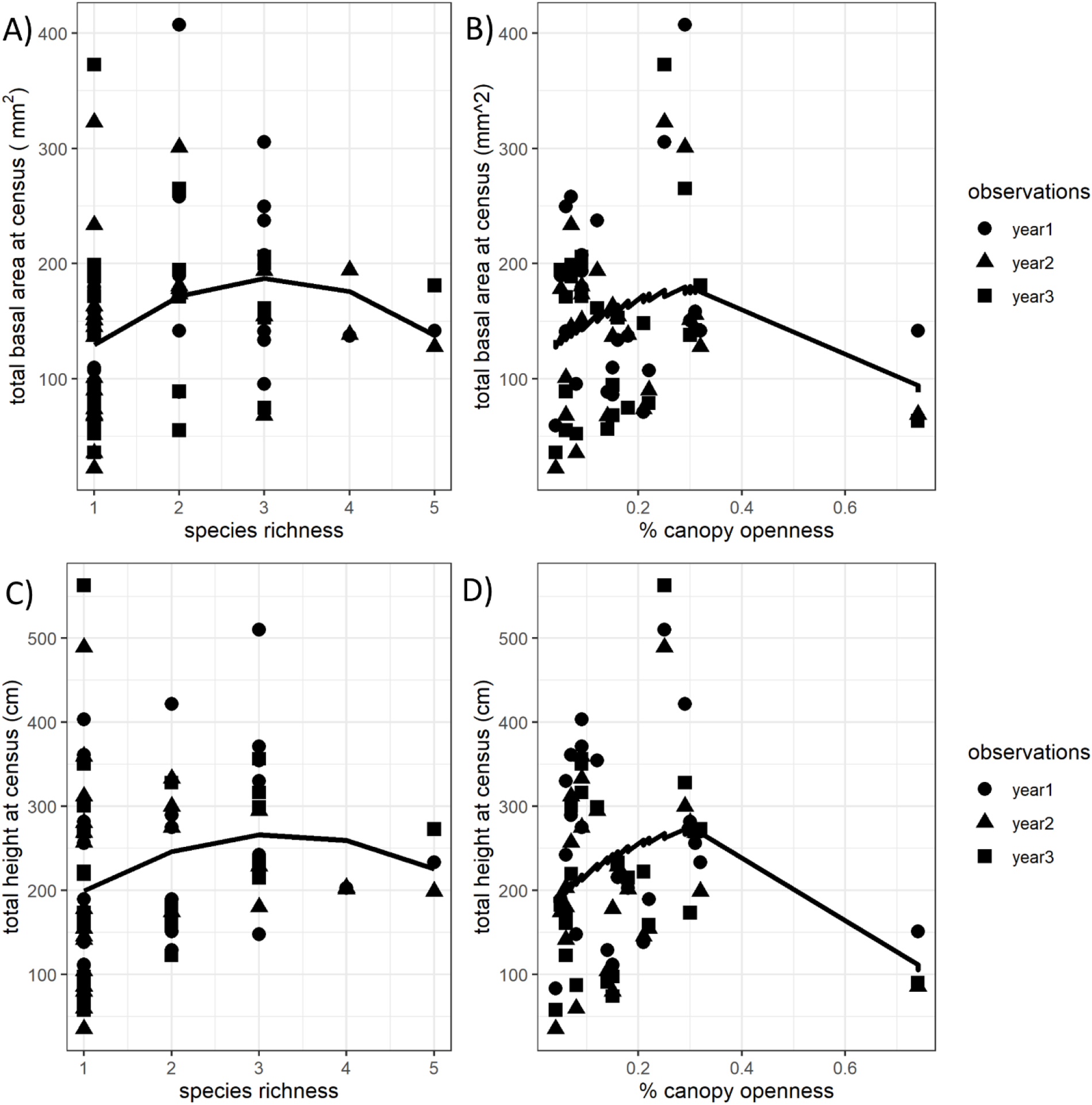
Observational data from long term plots. Generalised linear models of total basal area as a function of A) species richness and B) canopy openness and total height as a function of C) species richness and D) canopy openness.

## Discussion

We found that light modulates the positive effect of species diversity on biomass accumulation in seedling communities. In even-aged, experimental tree seedling communities, representing the onset of regeneration, community growth was increased by both light levels and species diversity. Further, we show that this increase in growth at the community level is mirrored at the individual seedling level as well, even across species varying in functional types and growth rates. Ultimately, we show that complementarity with non-random overyielding likely drives biomass accumulation in seedling communities under combined biotic and abiotic influences in seedling communities. Taken together, our results demonstrate a critical need to integrate biotic and abiotic factors that structure diversity and ecosystem functioning. Our results lay groundwork to further understanding of future forest diversity and ecosystem functioning by disentangling processes that structure tree communities under regeneration.

Our results show that in the first and most crucial year of growth, biomass accumulation in seedling communities can be influenced independently and additively by light and species diversity. Seedling communities planted with higher species richness and under canopies with higher light had higher values of basal area and height after a year of the experiment (Fig. 2) (Sapijanskas et al., 2013; Shen et al., 2021). This demonstrates a potential additive influence of diversity and light on community level seedling growth. However, the increase in biomass was clearly observable only at the higher level of species richness or in the deciduous canopy treatment indicating that growth responses to these factors are not linear at the community level (Fig 2). Lending weight to experimental results, such non-linear responses of plot-level biomass to light were also observed in the naturally regenerating communities; although light increased standing biomass of seedlings, a single plot with the most open canopy had low values of biomass (Fig 6). Although there were clear biomass gains with species richness in experimental plots, standing biomass increased weakly along the observed species diversity gradient (Fig 2, Fig 6). Under natural regeneration, species composition varies with microclimate conditions; for example, deciduous canopies with higher light have higher abundances of deciduous seedlings below them (Souza et al., 2014). Such niche partitioning by these functional groups and associated differences in growth rates could potentially explain weak relationships with species richness in natural communities.

We show that in seedling communities, overyielding with increasing richness can be explained by species complementarity rather than the increased probability of sampling faster growing or higher yielding species (Fig 3). Increasing diversity led to overyielding in communities (Fig 3), as expected from other studies on forest ecosystems (H. Pretzsch et al., 2015; Sapijanskas et al., 2013; Van de Peer et al., 2018). Beyond species richness, species functional composition influenced the survival and growth of communities (Fig S1, Fig S2) (Potvin & Gotelli, 2008; Salisbury & Potvin, 2015; Shen et al., 2020; Yang et al., 2013). However, despite considerable differences in growth rates among experimental species, inclusion of fast-growing species alone did not explain the observed increase in biomass in high-diversity treatments (Fig 3). In fact, selection effects decreased progressively from low to high species treatments, while complementarity effects increased (Fig 3). Such evidence about biodiversity-driven mechanisms in early life history stages of trees is lacking, but our results extend emerging evidence of complementarity in seedlings of forest species under constant environments (Bastias et al., 2021) and suggest that under heterogeneous environments, too, complementarity drives regeneration dynamics.

Deviating from purely competition-driven (complementarity) or probabilistic (selection) effects, our results indicate that the positive effect of biodiversity in heterogeneous environments can be driven by complementarity with non-random overyielding (Isbell et al., 2018). Contrary to expectations of differential responses by functional groups across different light conditions, individuals across species, on average gained more biomass with higher light and higher species richness in the community (Fig 4). In other words, fast-growing, high yielding, deciduous species continued to outperform slower growing evergreen species across all conditions, but species diversity and light increased yield for all species, leading to higher biomass in high light, high diversity treatments (Fig 5). Light and diversity-mediated increases in individual growth across species suggests strong resource and light competition within these communities, in agreement with other studies on tree seedlings (Lu et al., 2021; Madsen et al., 2020; Promis & Allen, 2017).

Grassland species as well as mature trees show significant spatial and temporal insurance effects, which we did not detect on tree seedlings (Bunker et al., 2005; Isbell et al., 2018; and as reviewed in Loreau et al., 2021). This implies that although functional differences in mature trees might stabilise production through differential growth under heterogenous environments, these differences do not produce similar asynchrony in seedling communities. Instead, strong complementarity in seedling communities is a possible consequence of high intraspecific competition in these stages or growth suppression by plant enemies (Inman-Narahari et al., 2016). Our initial results indicate the need to explore these mechanisms further across diverse ecosystems and life history stages.

Broadly, we show that the potential for diversity to maintain ecosystem functioning is affected by changing environmental conditions. In the highest light conditions in our experiment, biomass at the community level was not improved significantly or consistently with increasing diversity (Fig 2). Moreover, for individual seedlings, the proportional gain with increasing light was lower in high diversity treatments, indicating the limits of these positive outcomes, potentially due to other competitive interactions (Fig 4) (Rüger et al., 2011). These patterns may be driven by altered aboveground competition for light in high diversity communities, resulting from fast growth by some species and rapid vertical stratification observed during the experiment (Lu et al., 2021; Sapijanskas et al., 2014). Further, although light treatments in our study represent a strong driver of community structure in these forest understories, increased light could lead to both increased photosynthesis and increased water stress with consequent divergent effects on growth, that could interact with species diversity (O’Brien et al., 2017). While our results hold for both these processes acting together and represent realism to understand within-stand dynamics, these effects may be dissociated across forest stands and need further disentangling. More generally, multiple abiotic factors, including light, soil nutrients and soil moisture affect seedling dynamics in these forests. The strength of the biodiversity effect and the mechanisms through which it affects yield are likely to vary across forests structured by different environmental factors (García et al., 2018; Thompson et al., 2018, 2021).

Our findings have broad implications for understanding the interaction between multiple drivers and their influence on ecosystem functioning. Altered environmental conditions through global and local drivers combined with biodiversity loss across biomes have heightened the vulnerability of several ecosystem functions and services (Delzon et al., 2012; Heilpern et al., 2018). However, many mechanistic and simulation models that quantify forest futures continue to deal with single-species stands, simplified dynamics with light and focus largely on tree mortality than regeneration (Hans Pretzsch et al., 2015; Strigul et al., 2008). Our study adds to the understanding of future forest composition through by experimentally disentangling mechanisms that affect early growth stages (Shen et al., 2021; Uriarte et al., 2018). We show that biodiversity effects on biomass gain in early growing stages of long-lived tree species can be modulated by abiotic factors. In the light of renewed understanding from this study, predictions of future composition and ecosystem functioning need to include community-based approaches and regeneration dynamics. Updated models for regeneration that include interactions between biotic and abiotic drivers and outcomes at the community level are crucial for predictions of ecosystem functioning in a changing world.

## Acknowledgements

We acknowledge the support of the Department of Environment and Forest, Andaman and Nicobar Islands for conducting this work, under permit CCF/R&WP/9G(2)/Vol.XIV/741, and subsequent requests. This work was funded through a National Geographic Early Career Grant to KA. We are grateful to Ms. Nabanita Ganguly, Mr. Tarun Coomar, Ms. Deepa, Mr. Rashid and the staff at the Nayashahar Silviculture Nursery for permissions and logistical support. We also thank long-term field staff of the LEMoN plot in the Andamans – Anand James Tirkey, Sebian Horo and Ledu Kujur for their roles in data collection. This work was possible with community support at the Andaman Nicobar Environment Team. The authors would also like to thank Maria Uriarte for critical feedback during the design and analyses stages and Sarah Bruner and Pallavi Kache for writing support.

