## Supplementary table and figures for "Biodiversity effects on seedling biomass growth are modulated by light environment across functional groups"

Supplementary information

**Table S1:** Model selection for basal area and height in observed long-term dynamics plots.

| | | | | **Estimates** | | | |
|---|---|---|---|---|---|---|---|
| | | | Intercept | Linear term | Quadratic term | Random effect variance | AIC |
| Basal area | species richness | Linear | 118*** | 18.02 | | 1.50E-06 | 858.6 |
| | | Quadratic | 60.38 | 82.60* | -13.43* | 0 | **856.7** |
| | canopy openness | Linear | 150.10*** | 9.13 | | 1.79E-13 | 862.2 |
| | | Quadratic | 116.21*** | 363.55* | -538.12** | 46.03 | **858.6** |
| Height | species richness | Linear | 181.27 | 23.71 | | 1.76E-11 | 917.8 |
| | | Quadratic | 125.93 | 86.93 | -13.4 | 0 | 918.2 |
| | canopy openness | Linear | 233.21 | -43.69 | | 0 | 920.8 |
| | | Quadratic | 164.39*** | 641.40*** | -970.11*** | 69.15 | **913.9** |

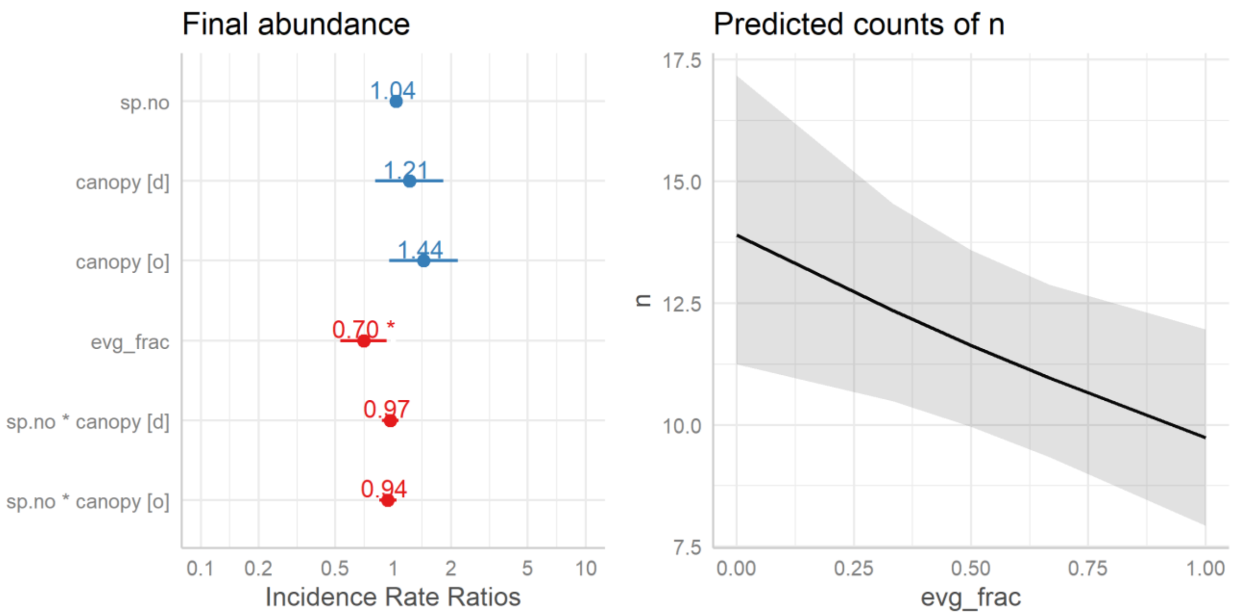


**Fig S1.** Generalized linear model with negative binomial family of errors modelling the number of individuals surviving at the end of 12 months of the experiment starting from equal abundance communities with 18 individuals each. Model deviance = 50.354, df=46, p=0.305.

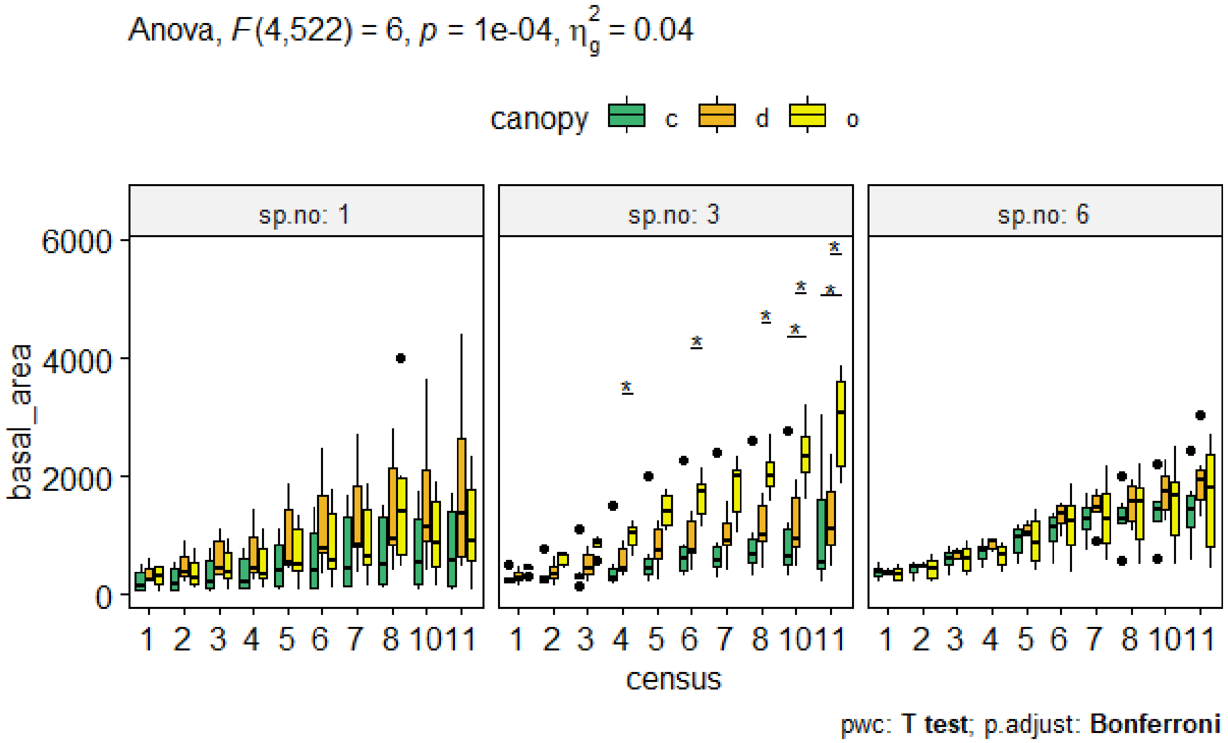


Fig S2. Repeated measures three-way ANOVA, F statistic = 6, p<0.001

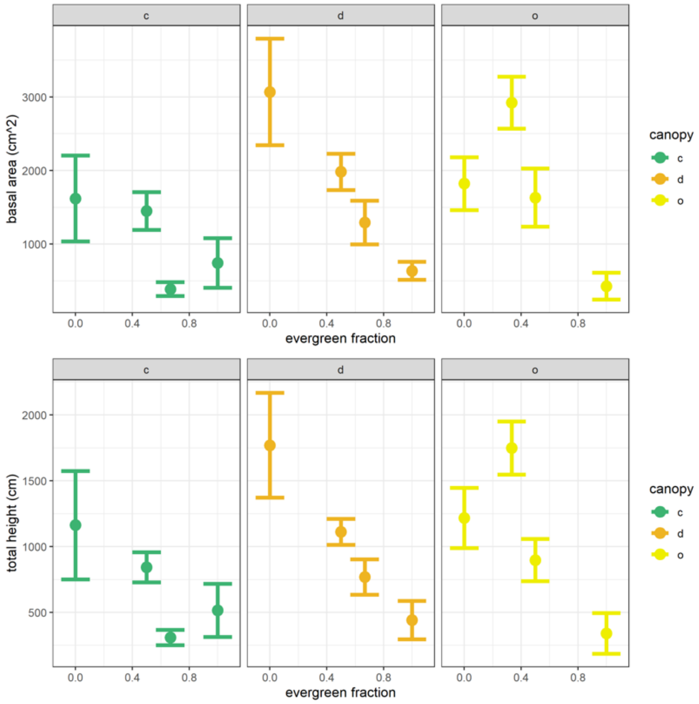


Figure S3: Final community properties as a function of functional composition. Evergreen fraction was calculated as the proportion of planted individuals from evergreen species in each community. Dots represent mean values at the end of the experiment across communities with similar functional composition. Error bars represent standard errors.
